# Dissecting the binding mechanisms of synaptic membrane adhesion complexes using a micropattern based cellular model

**DOI:** 10.1101/2024.03.17.584836

**Authors:** Nathalie Piette, Pierre-Olivier Strale, Matthieu Lagardère, Camille Saphy, Carsten Reissner, Matthieu Munier, Markus Missler, Ingrid Chamma, Matthieu Sainlos, Olivier Thoumine, Vincent Studer

**Affiliations:** Univ. Bordeaux, CNRS, Interdisciplinary Institute for Neuroscience, IINS, UMR 5297, F-33000 Bordeaux, France; Alveole, 30 rue de Campio Formio, 75013 Paris, France; Institute of Anatomy and Molecular Neurobiology, University of Munster, Germany

**Author notes:** equal contribution (Pierre-Olivier Strale, Matthieu Lagardère). equally contributing senior authors (Olivier Thoumine, Vincent Studer). **Author contributions:** N.P. optimized the protocol for the micropatterning, produced the micropatterned substrates, carried out all FRAP experiments, data analysis, made the figures, and contributed to the manuscript writing. P.O.S. and I.C. did the proof of concept for the micropatterning assay. M.L. and I.C. participated to initial FRAP experiments. M.L. contributed to the mathematical modeling of FRAP curves. C.S. performed single molecule pull-down experiments. C.R. and M.M. provided purified Nlg1-Fc and Nlg1-HaloTag proteins. M.S provided PSD-95 plasmids and helped with dye conjugation. O.T. and V.S. came up with the original idea, supervised the work, and wrote the manuscript. All authors reviewed the manuscript.

**Keywords:** Micropatterning, synaptic adhesion molecule, Diffusion, Binding

## Abstract

The formation of adhesive cell-cell contacts is based on the intrinsic binding properties between specific transmembrane ligand-receptor pairs. In neurons, synaptic adhesion molecules provide a physical linkage between pre- and post-synaptic compartments, but the strength and the dynamic of these complexes in their actual membrane environments remain essentially unknown. To access such information, we developed a versatile assay to measure the affinity and binding kinetics of synaptic ligand-receptor interactions, based on the immobilization of Fc-tagged ligands on micropatterned substrates combined with live imaging of fluorescently-tagged counter receptors in heterologous cells. We applied this strategy to study the heterophilic complex formed between neurexin-1β (Nrx1β) and neuroligin-1 (Nlg1), compared to the homophilic SynCAM1 complex. First, the control of ligand density combined to the measurement of steady-state receptor enrichment at micropatterns demonstrates the high specificity of the matching molecular interactions and allows for the quantification of the two-dimensional affinity of the interaction in a membrane environment. Second, long-term FRAP experiments performed on the two molecular complexes and fitted with analytical models, demonstrate a diffusion-limited regime for SynCAM1 and a reaction-limited regime for Nlg1. This analysis provides a very long bond lifetime of the Nrx1β-Nlg1 complex, which by comparison with a monomeric mutant of Nlg1, can be attributed to the constitutive dimerization of Nlg1. Finally, we used the stable Nrx1β-Nlg1 complex as a pseudo-synaptic platform to analyze the rapid binding kinetics between the scaffolding protein PSD-95 and the intracellular domain of Nlg1, dissecting the contribution of the different PDZ domains through the use of specific PSD-95 point mutants.

---

Adhesion is a fundamental process in cell biology, that regulates many processes including cell shape, migration, and differentiation, and is mediated at the molecular level by a wide range of membrane-associated ligand-receptor pairs. In neurons, specific families of adhesion molecules mediate developmental processes such as guidance and synaptogenesis. The most well-known synaptogenic proteins are pre-synaptic neurexins (Nrxs) and their post-synaptic binding partners neuroligins (Nlgs), as well as SynCAMs which can form both homophilic and heterophilic trans-synaptic interactions^*1–4*^. Those synaptogenic protein families contain multiple isoforms and splice variants, potentially generating thousands of combinations of binding partners ^*1*^. Moreover, genetic mutations in some of these molecules have been associated with autism spectrum disorders in humans, highlighting their importance in brain function ^*5*^.

Despite advances in the identification of such synaptic organizers, much remains to be learned about the multiple parallel or competing protein interactions that govern synapse assembly ^*6–8*^. In particular, a quantitative comparison of the affinity and kinetics of the various synaptogenic complexes may provide a better understanding of their specific biological role. While performing these measurements in living brain tissue is currently out of reach, several synaptic adhesion complexes have already been isolated *in vitro* and characterized by a range of biophysical methods ^*9–14*^. Although very informative, these techniques based on purified protein domains lack a critical membrane environment, and can lead to variable results, e.g. regarding the interaction rates of the Nrx1β-Nlg1 complex ^*10,14,15*^. In fact, the presence of the lipid bilayer which embeds those proteins in living cells can have profound impact on molecular diffusion and interactions, which crucially influences the clustering of receptors ^*16–18*^. Moreover, the reduced dimensionality imposed by the 2D membrane, the specific topography/curvature of the membrane, and the potential hindered receptor diffusion in contact zones, may overall introduce specific constraints to the ligand-receptor interaction rates that are not present when proteins are in solution ^*19,20*^.

Thus, there is a pressing need to engineer reductionist systems giving access to the quantification of interactions between specific ligand-receptor pairs in a proper membrane environment. Several cell culture methods allowing one to mimic synaptic ligand-receptor interactions have been developed in the past years, including co-culture and microsphere binding assays ^*21–27*^. However, these methods lack a control of the parameters governing these interactions (ligand density, geometry of the contact, receptor concentration) and thus prevent the proper quantitative analysis of specific adhesions. To circumvent these limitations, the micropatterning of proteins has recently been introduced to perform binding assays on living cells. In such assays, micron size patterns of ligands are printed on a glass cell culture substrate and fluorescently labelled counter-receptors are expressed in living cells ^*28–31*^. Such approaches have already led for example to the characterization of the dynamics and signalling properties of adhesion complexes in immune cells ^*32*^ and neurons ^*29*^.

Inspired by these earlier studies, we report here a biomimetic system in which recombinant Fc-tagged synaptogenic neuronal ligands are anchored to micro-patterned substrates and presented to heterologous COS-7 cells expressing GFP-tagged counter receptors, providing highly selective molecular recognition. Combining this system with Fluorescence Recovery After Photobleaching (FRAP) together with associated mathematical analysis in single cells, we characterized the kinetics of the different ligand-receptor complexes. By introducing well-chosen mutations in the receptor sequence, we also investigated the role of specific protein binding motifs in the adhesive interactions. We show that the Nrx1β-Nlg1 complex exhibits much slower binding kinetics than the SynCAM1 homophilic complex, most likely due to the dimerization of Nlg1. This property was further used to form a stable platform to dissect the intracellular binding kinetics of PSD-95 to Nlg1.

## Results

### Generating ligand-coated micropatterned substrates

To generate spatially controlled micropatterns onto which ligands can be coated, we exploited a technique previously engineered by ourselves, called Light Induced Molecular Adsorption of Proteins, or LIMAP ^*33*^. This method uses UV light focused through a microscope objective, to locally ablate polyethylene glycol (PEG) covalently bound to glass coverslips. Based on a predefined greyscale image imposed to a Digital Micromirror Device (DMD), we can create patterns of arbitrary geometry with micrometer resolution. By varying the UV dose, we can also directly control the grafted protein density. In practice, we patterned 5-µm wide stripes separated by 5-µm wide bands coated with PEG **(Fig. S1A)**. To ensure binding specificity and proper orientation of purified Fc-tagged synaptic adhesion proteins, i.e. Nrx1β-Fc and SynCAM1-Fc **(Fig. S1C)**, we used a robust chain of molecular interactions relying on the biotin-streptavidin pair (BSA-biotin, fluorophore-conjugated streptavidin, biotinylated anti-Fc antibody, and Fc-tagged ligand, sequentially) **(Fig. 1A)**. This amplification system is commonly designed to provide high ligand density and optimal protein orientation^*30,34*^. In this configuration, we measured the fluorescence intensity of streptavidin-AF647 (SA-AF647) and Fc-cy3 bound to the micropatterns at varying UV laser power (0-1000 mJ/mm^2^), and found that Fc-cy3 reached saturation at a power of 600 mJ/mm^2^, a value which was kept thereafter **(Fig. S1B-C)**. This corresponds to a density of ∼4500 molecules per µm^2^.

**Figure 1.**
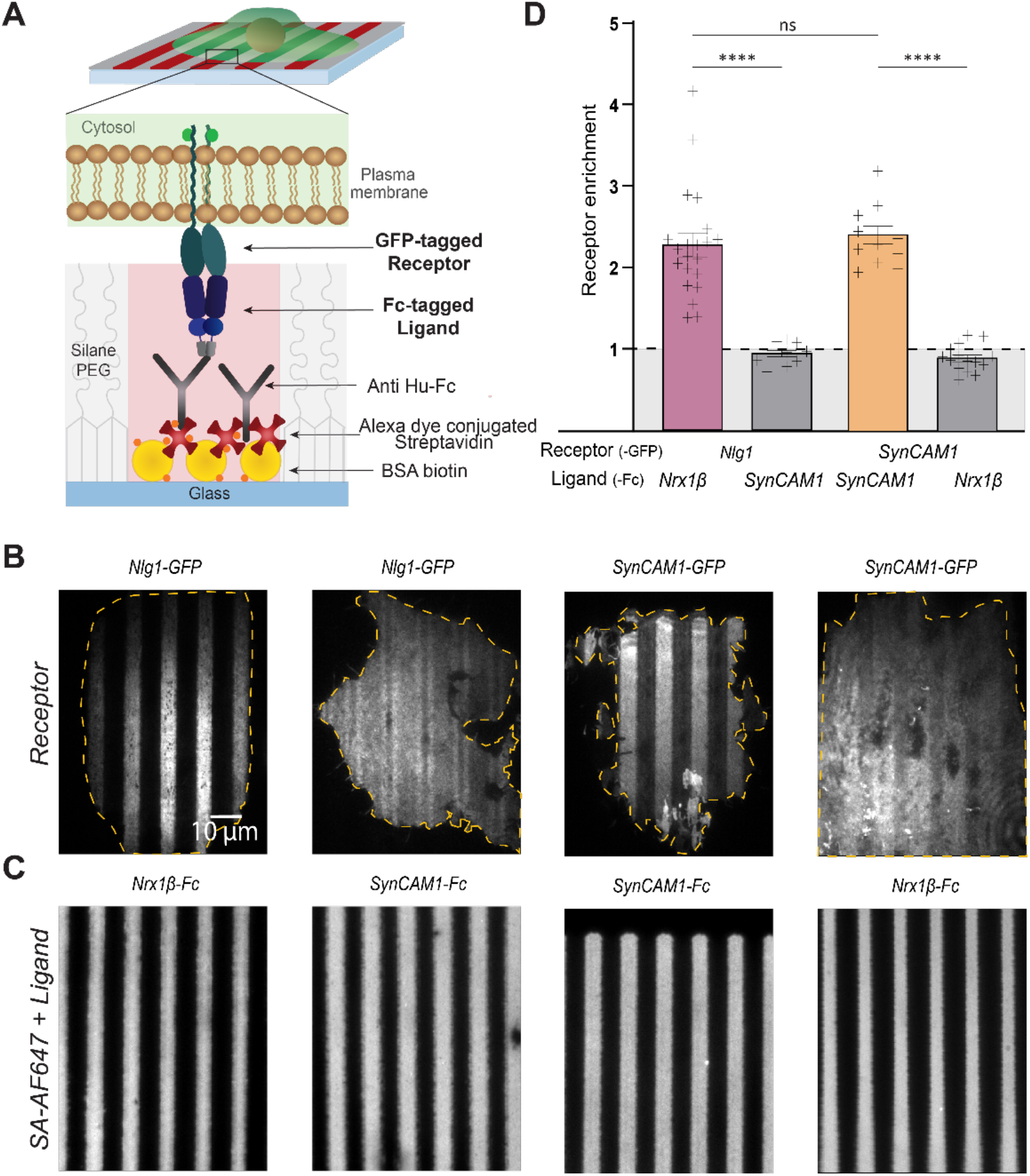
Micro-patterning assay to probe synaptic ligand-receptor interactions. (A) Schematic view of the assay, with a focus on the molecular interactions at the interface between the cell membrane and the micropatterned substrate. (B) Representative TIRF images of COS-7 cells expressing either Nlg1-GFP or SynCAM1-GFP and plated on micro-patterned lines coated with Nrx1β-Fc or SynCAM1-Fc. (C) Corresponding images of the streptavidin-A647 signal. (D) Enrichment factor of Nlg1-GFP or SynCAM1-GFP on Nrx1β-Fc or SynCAM1-Fc coated lines, respectively. Statistics are given as (number of cells/experiments): Nlg1-GFP on Nrx1β-Fc (21/3); Nlg1-GFP on SynCAM1-Fc (11/2); SynCAM1-GFP on SynCAM1-Fc (12/2); SynCAM1-GFP on Nrx1β-Fc (15/2). **** P < 0.0001; *** P=0.0002; ns>0.999 by kruskal-Walis test.

### Specific accumulation of cell-expressed receptors on ligand-coated micropatterns

We then used those micropatterned substrates to plate COS-7 cells electroporated with the putative counter-receptors bearing green fluorescent protein (GFP) tags, i.e. Nlg1-GFP and SynCAM1-GFP. To challenge the specificity of our model, we also performed the same assay with unpaired ligands and receptors. After several hours in culture, cells were observed by Total Internal Reflection Fluorescence (TIRF) microscopy **(Fig.1B)**, allowing us to monitor membrane receptors present in a thin optical section at the cell-substrate interface (< 100 nm) **(Fig. 1A)**. We quantified the receptor enrichment level (E), i.e. the ratio of the fluorescence intensity on the ligand-coated stripes vs PEG lines. The enrichment level was high for cells expressing Nlg1-GFP plated on Nrx1β-Fc (E = 2.3 ^+^/_-_ 0.15) and cells expressing SynCAM1-GFP plated on SynCAM1-Fc (E = 2.4 ^+^/_-_ 0.11) (**Fig. 1B,C)**. In contrast, no enrichment (E ≤ 1) was observed when cells were not expressing the appropriate counter receptor to the respective Fc-tagged ligand (e.g. Nlg1-GFP expressing cells on SynCAM1-Fc coated patterns, or SynCAM1-GFP on Nrx1β-Fc coated patterns), together revealing specific ligand-receptor recognition. In the absence of appropriate ligand-receptor pairing, cells still spread on the substrate most likely through non-specific adhesion to the pattern area and/or to the PEG, e.g. through the adsorption of serum proteins, allowing us to perform control measurements. Based on these results, we can experimentally correlate a high enrichment (E > 2) with a specific ligand-receptor binding interaction.

### Measuring affinities of ligand-receptor interactions in a cellular environment

We first exploited the steady-state values of receptor accumulation at ligand-coated micropatterns to estimate the intrinsic 2D dissociation constant between synaptic ligands and receptors. In a real biological system, knowing the free receptor and ligand concentrations is not straightforward. Here, the micropatterning system allowed us on one hand to control the ligand density [L_total_] in mol/µm^2^, and on the other hand to measure the corresponding accumulation of GFP-tagged receptors forming surface bound ligand-receptor complexes.

For a simple one-to-one ligand-receptor interaction, at thermodynamic equilibrium the rates of association and dissociation of the molecular complex compensate each other:

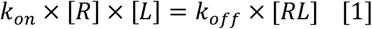

Where k_on_ is the association rate of the complex in mol/µm^2^/s, k_off_ the dissociation rate of the complex in s^-1^. [L] is the concentration in available surface bound ligand. Interestingly by measuring GFP fluorescence intensity values inside *I*_*ON*_ and outside *I*_*OUT*_ of the micropatterns, we have access to the surface membrane concentration of free receptors [R] out of patterns and bound receptors on the pattern [RL] in the same cell. We assume that we are in the linear fluorescence regime where the measured fluorescence signal is proportional to the local density of fluorescent molecules. In turn we can write:

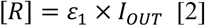

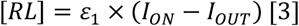

Where *ε*_1_ is the proportionality coefficient linking the fluorescence intensity measured on the camera to the local concentration of GFP molecules. We also assume and confirm (Supplementary figure S1 D) that in our experiments the density of Fc-tagged ligands is proportional to the intensity of the surface bound fluorescent streptavidin I_SA,_ via the proportionality coefficient *ε*_2_ .

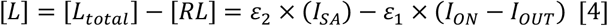

By combining equations [1] to [4], we can express the dissociation constant of the receptor-ligand pair K_d_=k_off_/k_on_ as a function of I_SA_, E = I_ON_/I_OUT_, and I_OUT_.

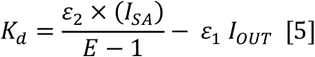

Which can also be written as:

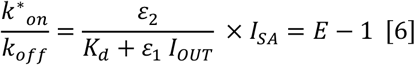

*where k*^*\**^_*on*_*= k*_*on*_ .[*L*] *is in units of s*^*-1*^

For each cell and for a given ligand-receptor pair, we expect a linear relationship between E-1 and I_SA_. To verify this, we took advantage of the grayscale patterning capability of our micropatterning system to tune the local density of immobilized Fc ligands so that each adhering cell can interact simultaneously with micro-patterns of different ligand density **(Fig. 2A-C)**. We then plotted E-1 as a function of I_SA_ for single cells and observe indeed the expected linear relation **(Fig. 2D)**.

**Figure 2.**
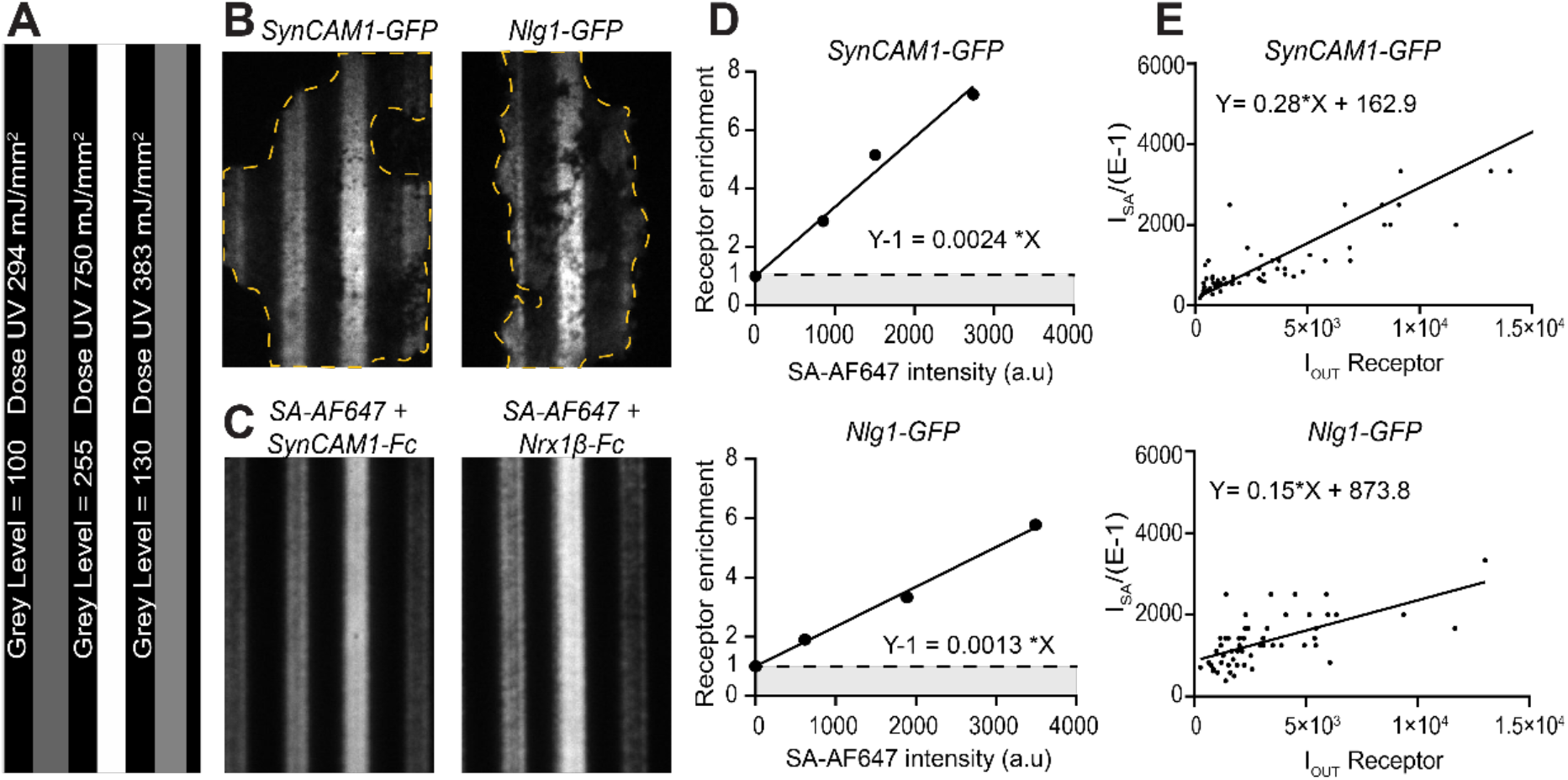
Dissociation constants computed from the surface density of ligands and receptors. (A) Part of an 8-bit grayscale projected image, comprising three different patterned lines (from left to right: 100 grey levels, width 24 pixels, 255 grey levels, width 20 pixels and 130 grey levels, width 22 pixels). (B) Representative examples of COS-7 cells expressing Nlg1-GFP or SynCAM1-GFP on micropatterned stripes coated with Nrx1β-Fc or SynCAM1-Fc, respectively. Three different ligand concentrations were printed using varying UV doses. (C) Corresponding streptavidin-AF647 signal on the stripes. (D) Graphs showing the relationship between Nlg1-GFP or SynCAM1-GFP receptor enrichment and the respective ligand density in one cell, expressed as the streptavidin-AF647 fluorescence level. (E) Graphs showing the relationship between ISA/(E-1) as a function of IOUT for all cells. Linear fitting gives the slope ε2/ε1 and the intercept at origin Kd/ε1. Statistics given as (number of cells/experiments): SynCAM1-GFP (59/1), Nlg1-GFP (54/2)

On top of that, electroporation gives rise to a wide range of receptor expression levels, as measured from the dispersion in the fluorescence intensity of the GFP reporter (especially true for SynCAM1-GFP). Interestingly, we can take advantage of this variability which leads to different concentrations of free receptors between cells. By rewriting again equation [5] we obtain:

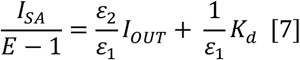

For a given receptor ligand pair, K_d_, *ε*_1_, *ε*_2_ should be equal so that each measurement of E and I_OUT_ for each single cell should fall on the affine relation [7]. We thus plotted for each ligand/receptor pair, 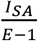 as a function of *I*_*OUT*_ for all cells. By a simple affine fitting we obtain K_d_/*ε*_1_ *and ε*_2_ */ε*_1_ **(Fig. 2E)**. In the case of SynCAM1, we obtain a nice agreement of our data with equation [7] (R^2^ =0.77). In the case of the Nrx1β-Nlg1interaction, the fit to the data is not good (R^2^ = 0.40). This can be explained by the fact that [7] holds only for a simple monomeric interaction which is not valid for the Nrx1β-Nlg1 complex. In addition, the expression level of for Nlg1 in COS-7 cells is lower than that of SynCAM1, which gives a lower range of variation.

Finally, we performed a titration assay with known dilutions of a purified GFP solution in the same TIRF imaging conditions as our cell experiments in order to extract *ε*_1_. In the case of synCAM1 we obtained a numerical estimation of the intrinsic two-dimensional K_d_=1,6.10^-22^ mol/µm^2^ (around 10 molecules/µm^2^). This measure done in a 2D membrane environment is hard to compare with bulk values from the litterature. Yet we show here how intrinsic affinities for a ligand/membrane receptor interaction can be obtained from multiple single cell micropattern measurements. Additionally, from equation [6] we show that the enrichment E in each cell is a measure of the reduced affinity 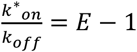 and how this measure depends on the expression level of the fluorescent receptor. Interestingly this reduced affinity can be measured by FRAP, which also allows the observation of the dynamics of ligand-receptor interactions, here again, in a membrane context.

### FRAP experiments reveal long-lasting bonds between Nlg1 and Nrx1β compared to SynCAM1

To gain insight into the dynamic processes that underlie the continuous binding and unbinding of synaptic membrane complexes, we performed Fluorescence Recovery After Photobleaching (FRAP) assays on our synaptogenic complexes. GFP-tagged synaptic receptors accumulated on their respective ligand-coated stripes were photo-bleached in less than a second on several areas of ∼3 µm diameter with intense and focused 491 nm laser light, then fluorescence recovery was monitored in TIRF mode for 5-20 min **(Fig. 3A-B)**. We expected a progressive replacement of bound photobleached receptors in micropatterned areas by freely moving unbleached receptors from nearby zones, that would reflect the turnover of ligand-receptor interactions. For Nlg1-GFP accumulated at stripes coated with Nrx1β-Fc, fluorescence recovery was biphasic, showing a rapid initial phase (< 2 min) up to 50%, followed by a much slower recovery phase up to 60% **(Fig. 3E)**. In contrast, the fluorescence recovery for SynCAM1-GFP accumulated at SynCAM1-Fc coated stripes was monophasic and more rapid (up to 80% in 5 min), indicating the formation of more labile bonds **(Fig. 3G)**. As a control, we used a Nlg1-GFP construct bearing two extracellular point mutations (L399A/N400A) that impair binding to Nrx1β ^35^. As expected, this mutant did not accumulate at Nrx1β-coated stripes (E ≤ 1), and showed very rapid fluorescence recovery (up to 100% in about 1 min), reflecting pure membrane diffusion **(Fig. 3F)**.

**Figure 3.**
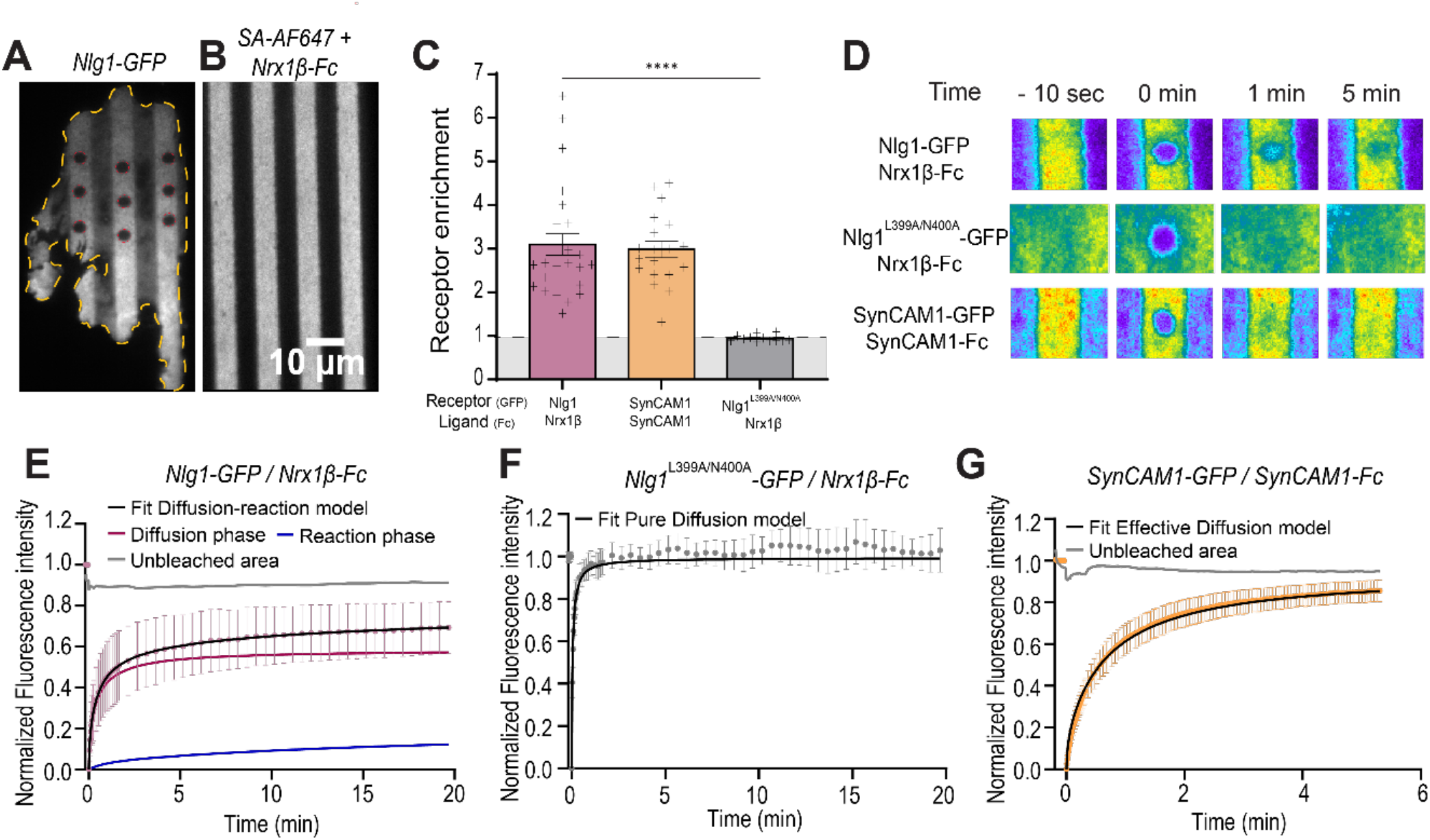
FRAP experiments on GFP-tagged SynCAM1 and Nlg1 accumulated at ligand-coated micropatterns. (A) Illustration of a COS-7 cell expressing Nlg1-GFP (left) spread on (B) micropatterned stripes coated with Nrx1β-Fc (streptavidin-A647 signal on right). Nine photobleached spots are shown at time zero. (C) Graph of GFP-tagged receptor enrichment on the micropatterned stripes in the different conditions **** P <0.0001 by Mann Whitney test. (D) Representative time lapse sequences of FRAP experiments performed on Nlg1^WT^-GFP or Nlg1^L399A/N400A^-GFP accumulated at Nrx1β-Fc coated stripes, or on SynCAM1-GFP accumulated at SynCAM1-Fc coated stripes. (E-G) Average recovery curves corresponding to the three conditions. The NLG1-GFP curve is fitted by the diffusion-reaction model, where the diffusion and reaction components are shown individually (red and blue curves, respectively). The Nlg1 ^L399A/N400A^-GFP curve was fitted by a pure diffusion model, while the SynCAM1-GFP curve was fitted with an effective diffusion model. Note the different time course for Nlg1-GFP and SynCAM1-GFP (20 min versus 5 min). The statistics are (number of cells/experiments): Nlg1^WT^-GFP (28/11); Nlg1^L399A/N400A^ -GFP (12/4); SynCAM1-GFP (19/4).

To quantitatively interpret our FRAP data, we used a comprehensive diffusion-reaction model described previously ^36,37^. The general solution is complex and solved only numerically, but it can be described by analytical functions in several simplified cases and can allow one to estimate the association k_on_^*^ (s^-1^) and dissociation k_off_ (s^-1^) constants:

1/ Pure diffusion when binding is negligible (k*_on_/ k_off_ <<1) with an equation [8] based on free diffusion (D_f_) with modified Bessel function ) of the first (I_0_) and second kind (I_1_) for a circular bleach area of radius w ^38^ .

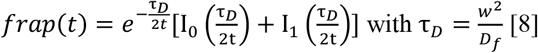

2/ Effective diffusion when reaction is fast compared to diffusion ((k^*^_on_ x w^2^/D_f_)>>1), the same equation [8] is used for this model but with an effective instead of a pure diffusion (τ_Deff_=w^2^/D_eff_). This apparent diffusion coefficient can be related to k^*^_on_/k_off_ but also, based on [6] to the enrichment E [9].

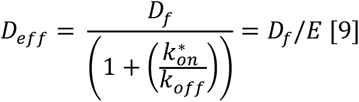

3/ Uncoupled diffusion and reaction when the adhesive interaction is slower compared to diffusion ((k^*^_on_ x w^2^ / D_f_) <<1). In this case multiple equations can be used, a full reaction dominant one when the diffusion can be completely ignored (Ceq=1(k_off_<<k^*^_on_, where Ceq is the equilibrium surface concentration of bound receptors) [10].

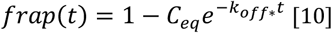

When the diffusion is present but occurs much faster and thus before the reaction (k^*^_on_ x w^2^/D_f_<<1), the FRAP recovery curve can be fitted [11] by be a combination of the equation [8] and [10].

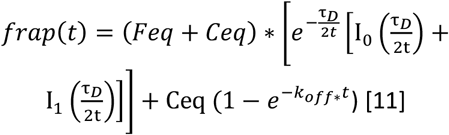

For each condition, we have determined the best fitting model. The parameters obtained from the fits are given in **Table 1**.

**Table 1.** Values extracted from fits of FRAP curves.

| Receptor | Ligand | Diffusion FRAP<br>($\mu\text{m}^2/\text{s}$) | Deffectif<br>($\mu\text{m}^2/\text{s}$) | $k_{\text{off}}$<br>($\text{s}^{-1}$) | Immobile Fraction % | Error FIT |
| --- | --- | --- | --- | --- | --- | --- |
| Nlg1<sup>WT</sup> | Nrx1β-Fc | 0.030 <sup>+</sup>/ - 0.005 | | 4.2.10<sup>-3</sup> <sup>+</sup>/ - 1.6.10<sup>-3</sup> | 22 <sup>+</sup>/ - 1.7 | 0.0052 |
| Nlg1<sup>E584A/L585A</sup> | Nrx1β-Fc | | 0.040<sup>+</sup>/0.026 | | 16 <sup>+</sup>/ - 1.47 | 0.035 |
| Nlg1<sup>L399A/N400A</sup> | Nrx1β-Fc | 0.18 <sup>+</sup>/ - 0.018 | | | 0.4 <sup>+</sup>/ - 0.3 | 0.26 |
| SynCAM-Fc | SynCAM1-Fc |  | 0.016 <sup>+</sup>/ - 0.0013 |  | 5 <sup>+</sup>/ - 0.8 | 0.055 |
| Nlg1 WT-HaloTag | Nrx1β-Fc | | | 1.5.10<sup>-3</sup> <sup>+</sup>/ - 0,8.10<sup>-3</sup> | 51 <sup>+</sup>/ - 13 | 0.0029 |

Values are presented at the mean ± SEM

In the FRAP analysis of the Nlg1^L399A/N400A^-GFP mutant ^39^ which does not adhere to Nrx1β-Fc patterns **(Fig. 3F)**, only pure diffusion is expected. The curve was nicely fitted using the pure diffusion model equation [8], yielding a diffusion coefficient (D) of 0.18 µm^2^/s. For the fluorescence recovery of Nlg1-GFP on Nrx1β-Fc stripes, characterized by a prolonged duration of FRAP (>30 min) and a distinctive curve shoulder, the uncoupled diffusion reaction model [11] was applied. The reaction component of the model, representing 19.5% (+/-1.5% SEM), enabled the extraction of a very low unbinding rate (k_off_ = 4.10^-3^ s^-1^), significantly smaller than values obtained through previous SPR experiments [14]. To confirm this notably slow turnover between Nlg1 receptors and Nrx1β-Fc ligand we used the same Nrx1β-Fc-coated micropatterned substrates but instead of cells expressing Nlg1-GFP we incubated a solution of purified dimeric Nlg1 C-terminally fused to Halo-tag and fluorescently-conjugated to AF488. Nlg1-Halotag-AF488 accumulated on Nrx1β-coated stripes similarly to Nlg1-GFP **(Fig. S2)**. Following photobleaching, fluorescence recovered gradually over time, missing the fast-initial regime corresponding to the 2D membrane diffusion of Nlg1 receptors. Here, the recovery stems from fluorescent Nlg1-Halotag molecules persisting in the surrounding solution and replacing photobleached molecules that gradually dissociate from their ligands. Consequently, the 3D diffusion is anticipated to be highly efficient and not limiting the recovery process. The FRAP curve was adequately fitted by a reaction-dominant model [10], yielding an unbinding rate of approximately k_off_ = 1.5.10^-3^ s^-1^, even slower than that observed in the presence of the cell membrane. Notably, a comparative analysis of the two conditions, when plotted side by side, reveals that the FRAP curve obtained for purified Nlg1-Halotag is similar to the long-term reaction regime observed for Nlg1-GFP. We also observe that the diffusion coefficient extracted from the diffusion phase of the Nlg1-GFP on Nrx1β-Fc curve (D = 0.030 µm^2^/s) is significantly lower than the diffusion coefficient measured with the Nlg1^L399A/N400A^-GFP mutant. This discrepancy can be attributed to two potentially intertwined factors: 1) The density of contacts established in the condition of interaction between Nrx1β and Nlg1^WT^ may contribute to a reduction in the diffusion coefficient, and 2) The observed system involves multiple states of interactions characterized by diverse association and dissociation rates. The diffusion phase observed may be governed by either effective diffusion or a reaction-dominant model. The latter possibility is plausible due to the constitutive dimeric nature of Nlg1^12,40,41^. The FRAP curve for SynCAM1-GFP on SynCAM1-Fc patterns fitted well with an effective diffusion model **(Fig. 3G)**. Interestingly, in each individual cell we were able to extract an enrichment E and an effective diffusion coefficient Deff from the FRAP curve fitting. The obtained values were compatible with equation [9] **(Fig.S3)**. By a linear fit of Deff vs 1/E we can extract the free diffusion coefficient of SynCAM1 1 in the contact area, Df = 0.041 µm^2^/s.

### Effect of constitutive Nlg1 dimerization on the interaction with Nrx1β-Fc

To study the impact of the dimerization of Nlg1, we generated a GFP-tagged Nlg1 construct bearing two point mutations (E584A/L585A) that impair Nlg1 dimerization ^24,42^ **(Fig. 4A)**. The monomerization of the Nlg1^E584A/L585A^-GFP has been controlled by measuring photobleaching steps by single molecule pull-down, with the dimer form of Nlg1-GFP and GFP-Nrx1 as a control **(Fig. S4)**. We observe that monomerization is not complete but 55% of Nlg1^E584A/L585A^ are in a monomeric form (showing 1 photobleaching step). We then analyzed the behavior of the corresponding proteins expressed in COS-7 cells at Nrx1β-Fc coated micropatterns by FRAP experiments. The Nlg1^E584A/L585A^-GFP mutant accumulated strongly (enrichment around 2.5), which is a little bit less than the enrichment of Nlg1^WT^-GFP on Nrx1β-Fc coated stripes. The FRAP curve showed a faster initial recovery (up to 75% in 2 min) and no further long-term increase in fluorescence after photobleaching **(Fig. 4C-D)**. This curve was well fitted by the effective diffusion model with D_eff_ = 0.04 µm^2^/s.z

**Figure 4.**
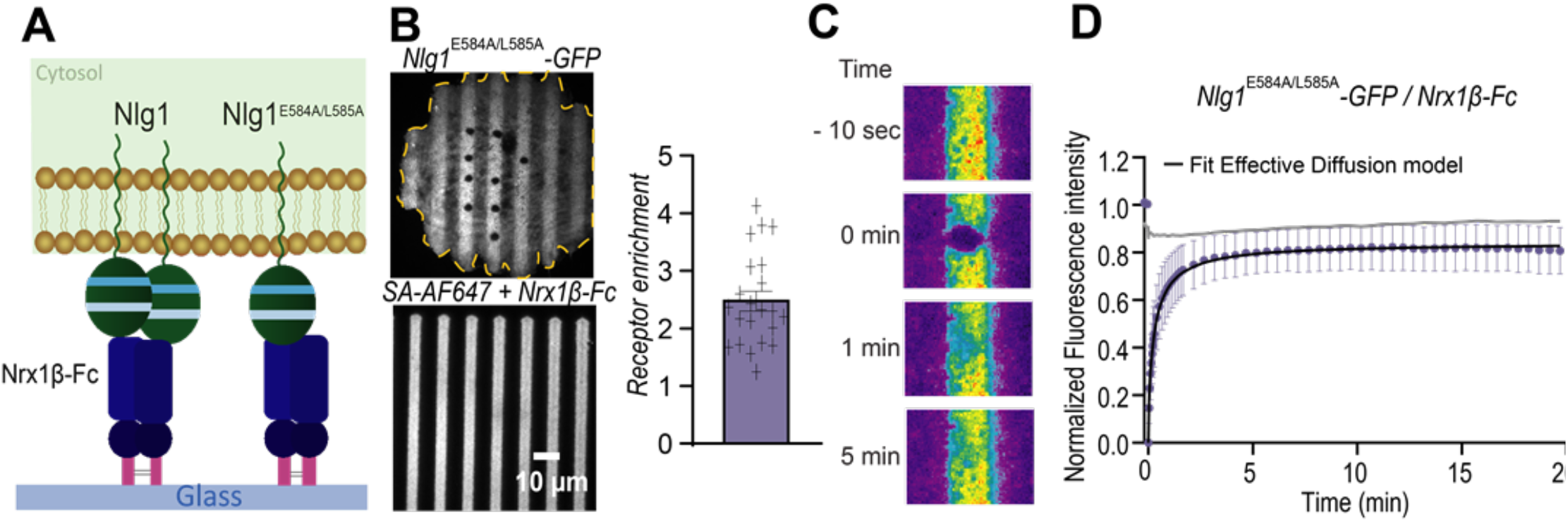
FRAP experiments on a Nlg1 dimerization mutant (A) Schematic representation of the dimeric and monomeric assays. (B) Illustration of a COS-7 cell expressing Nlg1^E584A/L585A^ -GFP (top) spread on micropatterned stripes coated with Nrx1β-Fc (streptavidin-A647 signal). Nine photobleached spots are shown at time zero. Graph of Nlg1^E584A/L585A^ -GFP enrichment on the micropatterned stripes; (C) Representative time lapse sequences and (D) corresponding recovery curves of FRAP experiments performed on Nlg1 ^E584A/L585A^ -GFP on Nrx1β-Fc coated substrates. Statistics given as (number of cells/experiments): (22/6)

Based on this observation, we fitted the rapid phase of the FRAP curve for dimeric Nlg1^WT^-GFP with the effective diffusion model [9]. We obtain an effective diffusion coefficient in the same range: D_eff_ = 0.03 µm^2^/s.

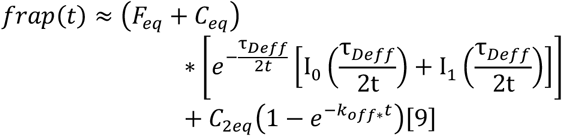

By fitting the FRAP curve with this equation, we can estimate that the proportion of Nlg1 molecules interacting simultaneously with two Nrx1β molecules should be around 10-20%.

In summary, Nlg1 appear to form rapid bonds with Nrx1β-Fc by mobilizing only one binding interface, but can also engage in long lasting bonds when the two lobes of the Nlg1 dimer are simultaneously binding two Nrx1β molecules, this configuration are probably favored by the high surface density of Nrx1β-Fc. When Nlg1 is forced to be in a monomeric form, as in the case of the Nlg1^E584A/L585A^ mutant, only the short lived bonds remain.

### Turnover of PSD-95 recruited at Nrx1β-Nlg1 adhesions

We demonstrated above the utility of our micropatterning assay to study interactions between transmembrane receptors and micropatterned ligands. Our next step was to characterize the interaction between a transmembrane protein recruited on the patterns and another intracellular membrane-associated partner. As a canonical molecular system, we have chosen PSD-95, a post-synaptic scaffolding protein anchored to the plasma membrane through a palmitoylation motif and which contains three PDZ domains enabling interactions with C-terminal motifs of various synaptic receptors, including Nlg1 ^26,43,44^. We co-electroporated COS-7 cells with recombinant GFP-tagged PSD-95 and AP-tagged Nlg1 and plated them on micropatterned substrates containing Nrx1β-Fc coated stripes **(Fig. 5A)**. Triggering the binding between membrane Nlg1 and Nrx1β-Fc caused the specific accumulation of PSD-95-GFP-WT at Nrx1β-Fc coated stripes **(Fig. 5B)**, with an enrichment factor E = 1.9 (^+^/_-_ 0.09 SEM), **(Fig. 5C)**. As a negative control, we used a Nlg1 mutant with a -5 aa C-terminal truncation (AP-Nlg1-Δ5) that removes the binding site to PSD-95 ^45,46^. The enrichment of PSD-95-GFP observed at Nrx1β-Fc coated stripes in COS-7 cells co-expressing Nlg1-Δ5 was close to 1.2 (^+^/_-_ 0.09 SEM) and significantly lower than that observed with Nlg1-WT, indicating that the Nlg1 PDZ-domain binding motif was involved in the recruitment of PSD-95. To further characterize the PDZ domains of PSD-95 involved in the binding to Nlg1, we used mutant PSD-95-GFP molecules carrying critical histidine to valine substitutions in PDZ domains 1, 2, or 3 (H130V, H225V, and H372V, respectively), that reduce the binding to C-terminal ligands ^47,48^. PSD-95-GFP carrying mutations in PDZ domains 1 and 2 (H130V/H225V) not expected to affect the binding to Nlg1, accumulated at Nrx1β-Fc coated lines as well as PSD-95^WT^-GFP (E = 2.5 ^+^/_-_ 0.2 SEM) **(Fig. 5B, C)**, indicating strong binding to Nlg1. Surprisingly PSD-95-GFP carrying a mutated PDZ domain 3 (H372V) expected to abolish the binding to Nlg1, also exhibited a relatively high PSD-95-GFP enrichment level at Nrx1β-Fc coated lines (E = 1.7 ^+^/_-_ 0.15 SEM), albeit significantly lower than the PSD-95^H130V/H225V^-GFP mutant. This result suggests that domains other than PDZ3 in PSD-95 might compensate for the binding to Nlg1.To quantify the interaction dynamics between Nlg1 and PSD-95, we then performed FRAP experiments on PSD-95^WT^-GFP or PSD-95-GFP mutants present at Nrx1β-Fc coated lines **(Fig. 5D**,**E)**. When co-expressed with full length AP-Nlg1, PSD-95^WT^-GFP showed a monophasic fluorescence recovery compatible with a diffusion-reaction model and reaching 80% in about 5 min, much quicker than the recovery of Nlg1-GFP on Nrx1β-Fc micropatterns. As a control, PSD-95^WT^-GFP co-expressed with HA-Nlg1^Δ5^ showed very fast fluorescence recovery that can be fitted with a pure diffusion model and yield a diffusion coefficient of 0.20 µm^2^/s, demonstrating that the Nlg1 C-terminal motif is responsible for stabilizing PSD-95 in this assay, as described in the literature^43^. When co-expressed with AP-Nlg1, the fluorescence recovery was barely faster for PSD-95^H372V^-GFP than for PSD-95^WT^-GFP **(Fig. 5E)**. Instead, the PSD-95^H130V/H225V^-GFP mutant showed a much lower recovery rate than PSD-95^WT^-GFP, confirming its higher enrichment level on Nrx1β-Fc patterns.

**Figure 5.**
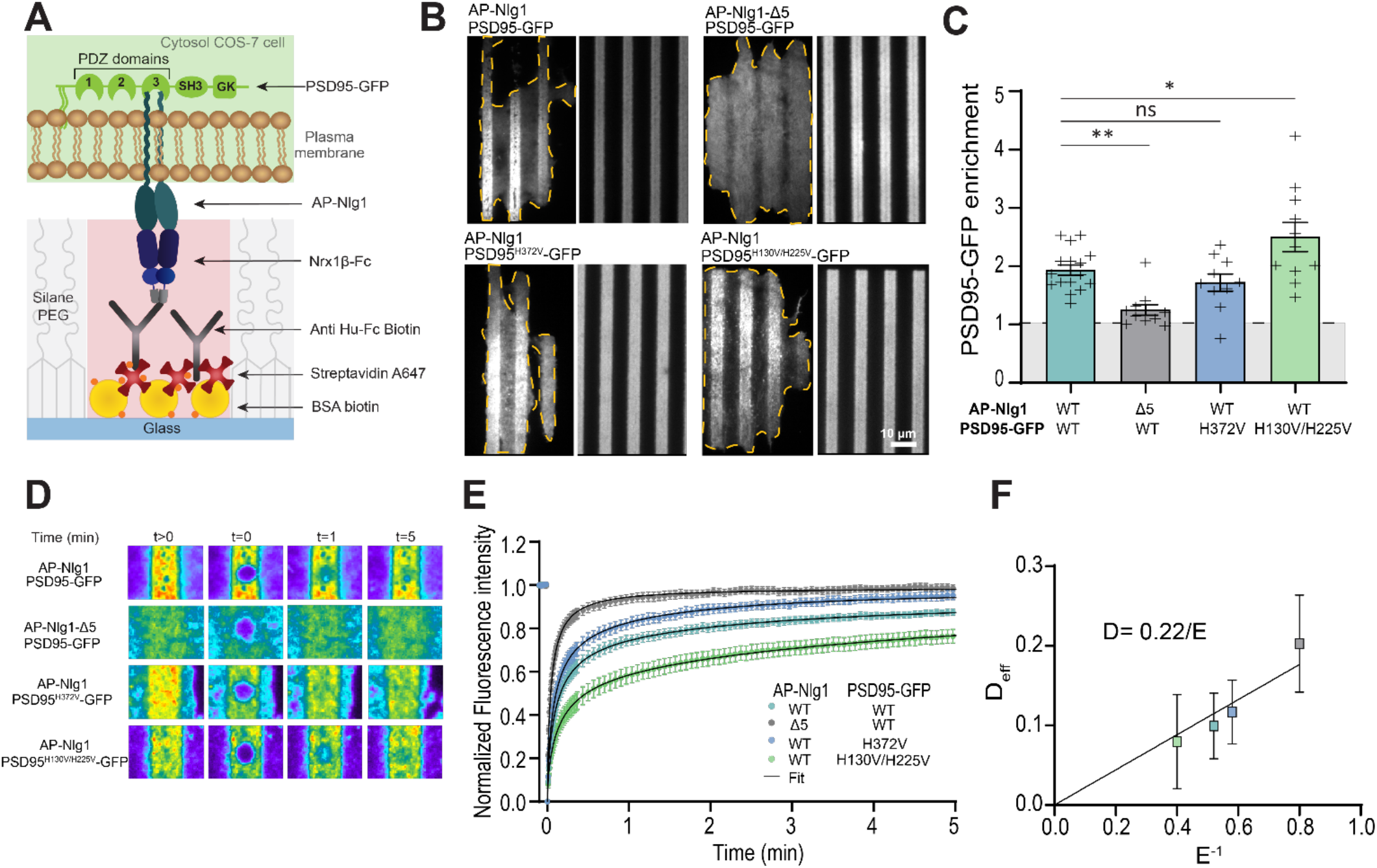
FRAP experiments on PSD-95-GFP at micropatterned Nrx1β-Nlg1 adhesions (A) Schematic diagram of the experiment. (B) Representative images of COS-7 cells co-expressing AP-Nlg1-WT or AP-Nlg1-Δ5 and PSD-95-GFP (either wild type or point mutants H130V/H225V or H372V) on micropatterned substrates coated with Nrx1β-Fc. PSD-95-GFP accumulation on Nrx1β-Fc coated lines in the different conditions. (C) Average PSD-95-GFP enrichment on Nrx1β-Fc coated lines versus regions between lines, in the different conditions (mean ± sem). ** P=0.0074, ns P= 0.1687, * P=0.0330 by one-way Anova multiple comparisons. (D) Representative FRAP sequences in the different conditions. (E) Average FRAP curves in the different conditions (mean ± sem). The curves were fitted by an effective diffusion model, except for PSD-95-GFP on Nlg1-Δ5 which was fitted by a pure diffusion model. (F) Graph showing the effective diffusion coefficient obtained from the FRAP curve fitting as a function of E^-1^ measured on the micropattern. Statistics given as (number of cells/experiments): WT (16/4), Δ5 (11/3), H372V (10/2) and H130V/H225V (11/2)

However, we cannot overlook the intrinsic turnover of Nlg1-Nrx1β adhesions at the substrate in this time frame. Indeed, by fitting the curve with the diffusion-reaction model we find that the slope of the reaction part gives a k_off_ between 5.10^-3^ s^-1^ and 8.10^-3^ s^-1^ depending on the condition, which is relatively close to the k_off_ obtained for the dimeric part of Nlg1 (4.10^-3^ s^-1^). Our hypothesis is that we are in the presence of a dual state model with a part corresponding to the PSD-95 interaction, which fits well with an effective diffusion model, and a part corresponding to the long dynamics of Nlg1/Nrx1β, reaction only [9]. This hypothesis is validated by the values of diffusion coefficient extracted from the first phase of the curve which are much higher for PSD-95^WT^ and the PSD-95 PDZ mutants compared to the Nlg1-Δ5 condition, where there is no interaction. Very interestingly, as expected theoretically from equation [**9]**, the effective diffusion coefficient obtained from the diffusive part of each FRAP curve fitting correlates nicely with the inverse of the enrichment measured on the micropatterns **(Fig. 5F)**. By a linear adjustment of D_eff_ vs E^-1^ we can obtain an estimation of the free diffusion coefficient of PSD-95-GFP at the membrane, D = 0.22 µm^2^/s. Together, these observations suggest that PDZ domains 1 and 2 compete with PDZ domain 3 for the binding to the Nlg1 C-terminal tail, and that when these domains are mutated, the interaction of Nlg1 with PDZ domain 3 is reinforced.

## Discussion

Accurate measurement of ligand-receptor affinity is of paramount importance for a detailed understanding of cellular signaling mechanisms. However, conventional methods often fall short in capturing the intricate dynamics within cellular environments. We propose here an approach that allows us to quantitatively assess ligand-receptor interactions at the single-cell level by analyzing the fluorescence intensity emitted by cell-surface receptors in contact with substrate regions micropatterned with their corresponding ligands. FRAP combined with appropriate mathematical models allowed for the estimation of the kinetics of the interactions in a membrane environment. The simple requisite of our system being to produce the Fc-tagged ligand and express its counter receptor in a cell line, our strategy should be applicable to many types of ligand-receptor pairs within different fields of biology, including development, immunology, and neuroscience. In fact, similar micropatterning approaches have already been used to characterize the signaling and dynamic properties of adhesion complexes such as those formed between EphB receptors and ephrins, or between T cell receptors and the intracellular kinase ZAP-70 ^32,49^.

On the technological side, the superposition of several layers of proteins on the substrate, i.e. BSA-biotin followed by streptavidin and biotinylated anti-Fc antibody to capture Fc-tagged ligands, provided a way not only to passivate the micropatterned area to decrease non-specific binding, but also to amplify ligand adsorption, together resulting in a highly reactive surface where only the appropriate counter-receptors accumulated. The technique should thus be expandable to screening purposes in multi-well plates, knowing that large protein banks of Fc-tagged ligands are already available ^50^. Using a tunable UV dose, the LIMAP technique also allowed us to vary ligand density on the micropatterns and study the effect on receptors within the same cell. This feature was instrumental to measure bidimensional binding affinities in single cell membranes. By performing a second round of printing with an orthogonal molecular chain (e.g. based on another protein tag) ^33^, this approach should also be applicable to the side-by-side patterning of different ligands, e.g. to study synergetic or competitive binding mechanisms between different ligand-receptor pairs.

On the biological side, we used this biomimetic model to dissect the interaction dynamics of canonical synaptic molecular complexes, namely the heterophilic extracellular binding between Nrx1β and Nlg1, the homophilic binding between SynCAM1 partners, and the intracellular binding between Nlg1 and PSD-95. Specifically, we showed that SynCAM1 forms relatively short-lived homophilic bonds as demonstrated by FRAP experiments, fitted well with an effective diffusion model. To our knowledge, there are very few reports in the literature focusing on the interaction kinetics between SynCAM family members, except one measurement of the dissociation constant between all the SynCAM family members ^51^. In this paper, the interaction SynCAM1/SynCAM1 was shown to be less labile than that computed from our results, i.e k_off_ = 0.0072 s^-1^. But it is difficult to compare 3D dissociation constants of proteins in solution to the 2D kinetic measurements performed here without a precise knowledge of the density of ligands and receptors ^19^.

In contrast, our data show that Nlg1 forms long-lived bonds with Nrx1β, mostly due to its dimeric nature. The introduction of point mutations in Nlg1 (L399A/N400A) shown to impair Nrx1β-Nlg1 binding *in vitro* ^39^ resulted in very little accumulation of the receptors and rapid FRAP kinetics on their respective ligands, confirming binding specificity. Moreover, a Nlg1 point mutant unable to dimerize (E584A/L585A) displayed a little less accumulation and more rapid FRAP kinetics on Nrx1β –Fc coated micropatterns than Nlg1^WT^, demonstrating that Nlg1 dimerization slows down its interaction with Nrx1β. This relatively slow kinetics is in agreement with that predicted from the gradual detachment of Nrx1β-Fc coated Quantum dots from primary neurons expressing Nlg1^42^. In comparison, SPR measurements performed on the Nrx1β –Nlg1 complex report disparate values, ranging from very fast kinetics when dimeric Nlg1 is adsorbed on the chip and interacts with soluble monomeric Nrx1β, i.e. k_off_ on the second time scale ^15^, to much slower kinetics when Nrx1β -Fc is grafted on the chip and interacts with dimeric soluble Nlg1, i.e. k_off_ = 0.015s^-1 14^. It is well known that Nrx1β-Fc also forms dimers thanks to two covalent disulfide bonds in the Fc domain. So, our first hypothesis was that the “dimerization” of Nrx1β by Fc domains affects the interaction. With the idea that monomeric Nrx1β can bind rapidly to either lobe of the Nlg1 dimer, but when arranged in dimeric form (Nrx1β-Fc) establishes long-lasting bonds with the Nlg1 dimer. But more recent measurements by isothermal calorimetric titration, where the molecules are in soluble form, describe an interaction with a strong exothermic reaction and a K_d_ of 390 nM between the ectodomain of Nlg1 and monomeric Nrx1β^52^. In the same article, measurements carried out in SPR with soluble monomeric Nrx1β give a K_d_ twice as high, with a value of 718 nM^52^. This measurement was performed in the presence of dimerized Nrx1β via Fc. The hypothesis of the creation of a double state due to Fc therefore does not appear to be the cause of the measurement differences described in previous articles^14,15^, but really to the dimeric form of Nlg1 and some rebinding effect promoted by the high density of Nrx1β.

This strong binding interface between Nlg1 and Nrx1β-Fc was further used as a template to explore the interaction dynamics between the Nlg1 C-terminus and PSD-95, revealing a complex interplay between the 3 PDZ domains. A potential mechanism to explain our findings would be that one C-terminal tail of a Nlg1 dimer interacts preferentially with PDZ domain 3 of PSD-95, while the other C-terminal tail binds PDZ domains 1 or 2 on the same PSD-95 molecule. When PDZ domains 1 and 2 are mutated, the Nlg1 dimer interacts (more strongly) which two PDZ domains 3 on neighboring or dimerized PSD-95 molecules ^53^.

These results obtained in heterologous cells should be valuable to better understand the dynamics of synaptic complexes in neurons. For example, SynCAM1 forms small domains at the periphery of synapses that move away upon an NMDA treatment which mimics long term depression (LTD) ^54^. The fast kinetics of the homophilic SynCAM1 complex reported here is compatible with a rapid regulation of its adhesiveness at the synaptic belt. Nrx1 and Nlg1 were found to form stable sub-micron clusters facing each other at the core of the synaptic cleft ^55–57^. The stability of these trans-synaptic complexes shown in our biomimetic assay may explain the fact that neurons are endowed with the ability to proteolytically cleave Nlg1 below its extracellular adhesive domain, when synapses are weakened in response to chemical LTD ^58–60^. Dimerization of Nlg1 is required for the ability of Nlg1 to induce new synapses in a co-culture assay, and to increase AMPA receptor-dependent synaptic transmission in organotypic hippocampal slices, these effects being mediated by the clustering of pre-synaptic neurexins ^24,61^. Our results showing that the monomeric mutant Nlg1^E584A/L585A^ unbinds quickly from with Nrx1β supports the concept that long-lasting trans-synaptic interactions between dimeric Nlg1 and Nrxs are necessary to mediate the full function of Nlg1 at synapses. Whether Nrxs can also form dimers through interactions with multimodal pre-synaptic proteins such as CASK ^62,63^ remains unclear, but this would represent an additional mechanism to reinforce trans-synaptic bonds with Nlgs. In fact, FRAP experiments performed on synaptic Nlg1 in neurons reveal a longer turnover rate than the one measured here on Nlg1 accumulated at Nrx1β-Fc coated micropatterns ^55,64^ suggesting that, in addition to pre-synaptic Nrx1, other molecular interactions stabilize Nlg1 at synapses. Indeed, Nlg1 can interact with a variety of intracellular partners, including PDZ domain containing scaffolding proteins such as PSD-95 and the WAVE regulatory complex ^26,29,65^. Interestingly, FRAP experiments performed on PSD-95-GFP accumulated at Nlg1 clusters induced by Nrx1β-Fc cross-linking in neurons ^26^ reveal a much lower turnover rate than the one observed here for PSD-95-GFP expressed in COS-7 cells and accumulated at Nrx1β-Fc coated micropatterns. These results suggest a bi-directional stabilization of Nlg1 and PSD-95 in neurons, most likely through the formation of multivalent bonds that are dominant at synapses. Further modeling work is necessary to predict the actual turnover rate of such proteins at synapses based on the individual protein-protein kinetic rates measured here.

## Acknowledgements

We thank T. Biederer, P. Scheiffele, S. Okabe, and A. Ting for the generous gift of plasmids, B. Tessier and S. Benquet for molecular biology; R. Sterling, A. Castets, and the cell culture facility of the Institute; J.

This work received funding from the Fondation pour la Recherche Médicale (“Equipe FRM” DEQ20160334916), French Ministry of Research, Agence Nationale de la Recherche (grant « Synthesyn » ANR-17-CE16-0028-01), Investissements d’Avenir Labex BRAIN (« Single Pull » ANR-10-LABX-43), the national infrastructure France BioImaging (grant ANR-10INBS-04-01), the Conseil Régional Aquitaine (« SiMoDyn ») and Deutsche Forschungsgemeinschaft SFB1348 - TP A03.

## Materials and methods

### DNA plasmids

SynCAM1-GFP (+420) ^66^ was kind gift from T. Biederer (Yale University, New Haven, USA). GFP-Nrx1β ^67^ was kind gift from M. Missler (Münster University, Germany). HA-Nlg1-GFP (-21) ^21^ and AP-Nlg1 ^68^, and were kind gifts from P. Scheiffele (Biozentrum, Basel, Switzerland) and A. Ting (Stanford University, Palo Alto, CA, USA) respectively. The PSD-95-GFP (+253) plasmid and the H130V-H225V and H372V mutant versions of this plasmid were kindly offered by M. Sainlos (IINS, Bordeaux), these plasmids have already been described ^69^. AP-Nlg1Δ5 was already described^70^. All Nlg1 DNA constructs contained both A and B splice inserts. HA-Nlg1^E584A/L585A^-GFP (-21) was generated by sub-cloning the mutated part of Nlg1 from HA-Nlg1^E584A/L585A^, already described ^71^, in HA-Nlg1-GFP (-21) at EcoRV/KpnI sites. The double mutation L399A/N400A was inserted in the HA-Nlg1-GFP (-21) by replacing the EcoNI-EcoRV WT sequence of Nlg1 by the insert generated by Eurofins containing the mutations.

### Purified proteins

Nrx1β and SynCAM1 C-terminally fused to human Fc were produced by transfecting HEK cells with the corresponding plasmids containing a hygromycin resistance, and culturing the cells for several weeks under hygromycin pressure (0,5 mg/ml) in serum-free medium. 500 mL of conditioned medium was collected in total and frozen at - 20°C. The Fc-tagged proteins were affinity-purified on a protein G column, and eluted with glycine, as described ^72^. Nlg1-HaloTag was generously provided by Carsten Reissner. The coupling with the Halotag-Alexa488 ligand (Promega, G1002) to Nlg1-Halotag was performed by incubating Nlg1-HaloTag with an excess (5X) of Halotag-AF488 ligand at 4°C under agitation overnight. Subsequently, the solution was passed through a centricon (10 kDa) to remove the excess free ligand. The Nlg1-Halo-AF488 was resuspended to a concentration of 0.5 mg/ml in a solution containing 10 mM Hepes and 300 mM NaCl. Aliquots of the solution were stored at -20°C after filtration. To examine the purity of the produced proteins, we ran samples of 1 µg of each protein on SDS-PAGE and immunoblotted the proteins using anti-human Fc antibodies.

Bovine serum albumin biotinylated was purchased from Thermo fisher. Cy3-conjugated Human Fc fragment and Biotin-SP-conjugated Goat Anti Human IgG Fc was purchased from Jackson Immunoresearch. Streptavidin, Alexa Fluor 405 or 647 conjugates were purchased from molecular probes.

### Glass micropatterning with Fc-tagged synaptic proteins

Substrate micropatterning was performed in a clean room. 18 x 18 mm High Precision Microscope Cover Glasses (Marienfeld SUPERIOR Germany Cat.No. 0107032) were fist activated by a 5 min air plasma treatment (Harrick Plasma, Ithaca, NY, USA). An 18 mm, 250 µm thick piece of Poly(dimethylsiloxane) (PDMS), with one square hole was cut in a transparent silicone film (BISCO HT-6240, Rogers Corporation Carol Stream, IL, USE) with a craft Robo Pro vinyl cutter (Graphtec, Irvine, CA, USA) and placed on the coverslip. Then the coverslips are placed in a closed chamber with 150 µl of (3-aminopropyl) triethoxysilane (Sigma-Aldrich, Saint-Quentin Fallavier, France) and a desiccator (Sigma-Aldrich) for 1hr. A solution of poly (ethylene glycol) -Succinimidyl Valerate, MW 5000 (Peg-SVA, Laysan Bio, Arab, AL, USA) at 228.6 mg / ml in 10 mM carbonate buffer pH> 8, is incubated in the well for 1 h and rinsed thrice with phosphate buffered saline (PBS) without drying the surface. Micropatterning was achieved using the PRIMO system coupled to the Leonardo software (Alveole, France) and mounted on an inverted microscope (Nikon Eclipse Ti-E, Japan). The pattern chosen is a series of 5 µm-wide parallel lines separated by 5 µm. After alignment of the well center with the UV pattern by moving the scanning stage, a solution of photo-initiator (PLPP, Alveole, France) was incubated on the sample. The UV-activated photo-initiator molecules locally cleave the PEG chains in a UV dose dependent manner and allows for protein adsorption to the UV-exposed areas. After rinsing the PLPP solution, the well was incubated with biotinylated bovine serum albumin (BSA) diluted to 100 µg/ml in PBS. Successive incubations were made with streptavidin-AF647 (FRAP), biotinylated anti-Fc antibody, and Purified-Fc protein (all at 50 µg/mL in PBS) all the incubations are done at RT except for Purified-Fc protein where the incubation is done at 4°C. The BSA-biotin and the streptavidin are incubated 3 min, the biotinylated anti-Fc antibody 10min and the Purified-Fc protein overnight. Between proteins, three washes were made in PBS without drying the surface.

### Cell culture and electroporation

COS-7 cells (ATCC) were cultured in FluoroBrite Dulbecco’s modified Eagle’s medium (DMEM; GIBCO/BRL) supplemented with 10% fetal bovine serum (FBS), 100 units mL^-1^ penicillin, 100 µg.mL^-1^ streptomycin, 1% Glutamax and 1% Sodium-Pyruvate in a 37°C-5% CO_2_ atmosphere. two days before the experiments, cells were rinsed twice in warm PBS, trypsinized for 5 min, mixed in culture medium, and centrifuged for 5 min at 1,000 rpm. The pellet containing around 1M cells was suspended in 100 µL electroporation medium, placed in a cuvette, and electroporated with 1-3 µg of each of the following plasmids, using the Amaxa Nucleofector system (Lonza): and GFP-Nrx1β, Nlg1-GFP or SynCAM1-GFP (for FRAP). For FRAP experiments on PSD-95-GFP, cells were co-electroporated with AP-NLG1 or AP-NLG1Δ5, together with PSD-95-GFP (either WT or point mutants H130V/H225V or H372V), in a 4:1 NLG1:PSD-95 ratio. Electroporated cells were resuspended in culture medium, plated in 12-well plates and placed back in the incubator for 36 hr. Then, cells were detached with trypsin and seeded on micropatterned glass coverslips at a concentration of 20,000 cells per coverslip, after 2 hours of incubation, rinsed to remove non-adherent cells and imaged 1-3 hrs later.

### FRAP experiments and analysis

COS-7 cells expressing the different GFP-tagged proteins (Nlg1, SynCAM1, or PSD-95) were seeded for 3 hrs on Fc-tagged ligand coated micropatterned substrates in culture medium, before being mounted in Tyrode solution in an Inox chamber. Cells were observed on a Nikon TiE Eclipse inverted microscope through a 100x/1.49 oil immersion objective, using 491 nm laser light under Total Internal Reflection Fluorescence (TIRF). Enrichment was quantified as the GFP level measured on Ligand-Fc coated lines divided by the level between those lines. The laser bench used for TIRF illumination has a second optical fiber output connected to an illumination device containing two x/y galvanometric scanning mirrors (ILAS, Roper Instrument) steered by MetaMorph. It allows precise spatial and temporal control of the focused laser beam at any user-selected region of interest within the sample for targeted photobleaching. Switching between the two fibers for alternating between imaging and bleaching is performed in the millisecond range using an AOTF. Oblique illumination was performed using the 491 nm laser at low power (300 µW at the front of the objective) to image molecules in the plasma membrane close to the substrate plane. After acquiring a 10 sec baseline at a 0.5 Hz frame rate, rapid and selective photo-bleaching of 9 circular areas of diameter 10 pixels was achieved at high laser power (3mW at the front of the objective), during 560 ms. Fluorescence recovery was then recorded immediately after the bleach sequence for 5 to 20 min at a 0.5-1 Hz frame rate. Observational photobleaching was kept very low, as assessed by measuring control unbleached areas nearby. FRAP curves were obtained by computing the average intensity in the photobleached area, after background subtraction, normalization with a none photobleach region to compensate bleaching if it is necessary and normalized between 1 (baseline) and 0 (time zero after photo-bleaching).

### Single molecule pull-down

Single molecule pull-down was performed essentially as described^73^. Briefly, cleaned glass coverslips (VWR) were treated with N-(2-aminoethyl)-3-aminopropyltrimethoxysilane (United Chemical Technologies, #A0700), then incubated with mPEG-succinimidyl valerate (SVA) containing a 1:100 fraction of biotin PEG-SVA (both from Laysan Bio MW 5,000 Da). Coverslips were dried and stored at -20°C. Just before the experiment, coverslips were incubated with 1 µM NeutrAvidin for 5 min (Invitrogen, A2666), followed by 10 nM biotinylated anti-GFP antibody for 15 min (ABCAM, 6658), both diluted in T50 buffer (10 mM Tris-HCl, 50 mM NaCl, pH = 7.5). Two days before the experiment, HEK cells were transfected with GFP-Nrx1β, Nlg1-GFP, or Nlg1E584A/L585A-GFP using Lipofectamine (Invitrogen) in optiMEM buffer. Two hours before the experiment, cells were detached in Ca-free PBS (30 min, 37°C), then centrifuged at 3,000 rpm for 2 min. The cell pellet was dissolved in 100 μL lysis buffer containing 150 mM NaCl, 10 mM Tris-HCl, 1 mM EDTA, 1 % Igepal, and protease inhibitor cocktail (Pierce). Samples were rotated for 2 hrs at 4°C to solubilize proteins, then centrifuged at 14,000 rpm for 15 min to remove aggregates. The supernatant was kept on ice and diluted 1:200 in observation solution (135 mM NaCl, 5 mM KCl, 2 mM CaCl2, 1 mM MgCl2, 10 mM HEPES, 4 mM Trolox, 40 mM D-glucose, 0.03% Igepal, pH = 7.4). 100 µL of this mix was added to glass substrates for several minutes, then rinsed out with fresh solution when the molecular density reached about 50 per field of view (256 x 256 pixels of 160 nm each = 41 x 41 µm). Single GFP-Nrx1β molecules were observed with a 491 nm laser under TIRF illumination on the same microscope used for SPT-PALM (Nikon TiE Eclipse equipped with the 100X/1.49 N.A. oil immersion objective). GFP-tagged molecules did not stick to a control coverslip containing no secondary anti-GFP antibody, revealing binding specificity to the GFP tag. Ten acquisition sequences of 700 frames for each condition were recorded at 30 Hz on the EMCCD camera (Evolve, Roper Scientific, Evry, France), resulting in the photobleaching of nearly all immobilized GFP-tagged molecules. Images were analyzed using Metamorph by measuring the average fluorescence intensity in small regions drawn around individual molecules, and the number of photobleaching steps was determined visually for more than 120 molecules per condition.

## Author contributions

N.P. optimized the protocol for the micropatterning, produced the micropatterned substrates, carried out all FRAP experiments, data analysis, made the figures, and contributed to the manuscript writing. P.O.S. and I.C. did the proof of concept for the micropatterning assay. M.L. and I.C. participated to initial FRAP experiments. M.L. contributed to the mathematical modeling of FRAP curves. C.S. performed single molecule pull-down experiments. C.R. and M.M. provided purified Nlg1-Fc and Nlg1-HaloTag proteins. M.S provided PSD-95 plasmids and helped with dye conjugation. O.T. and V.S. came up with the original idea, supervised the work, and wrote the manuscript. All authors reviewed the manuscript.

## Supplemental figures

**Figure S1.**
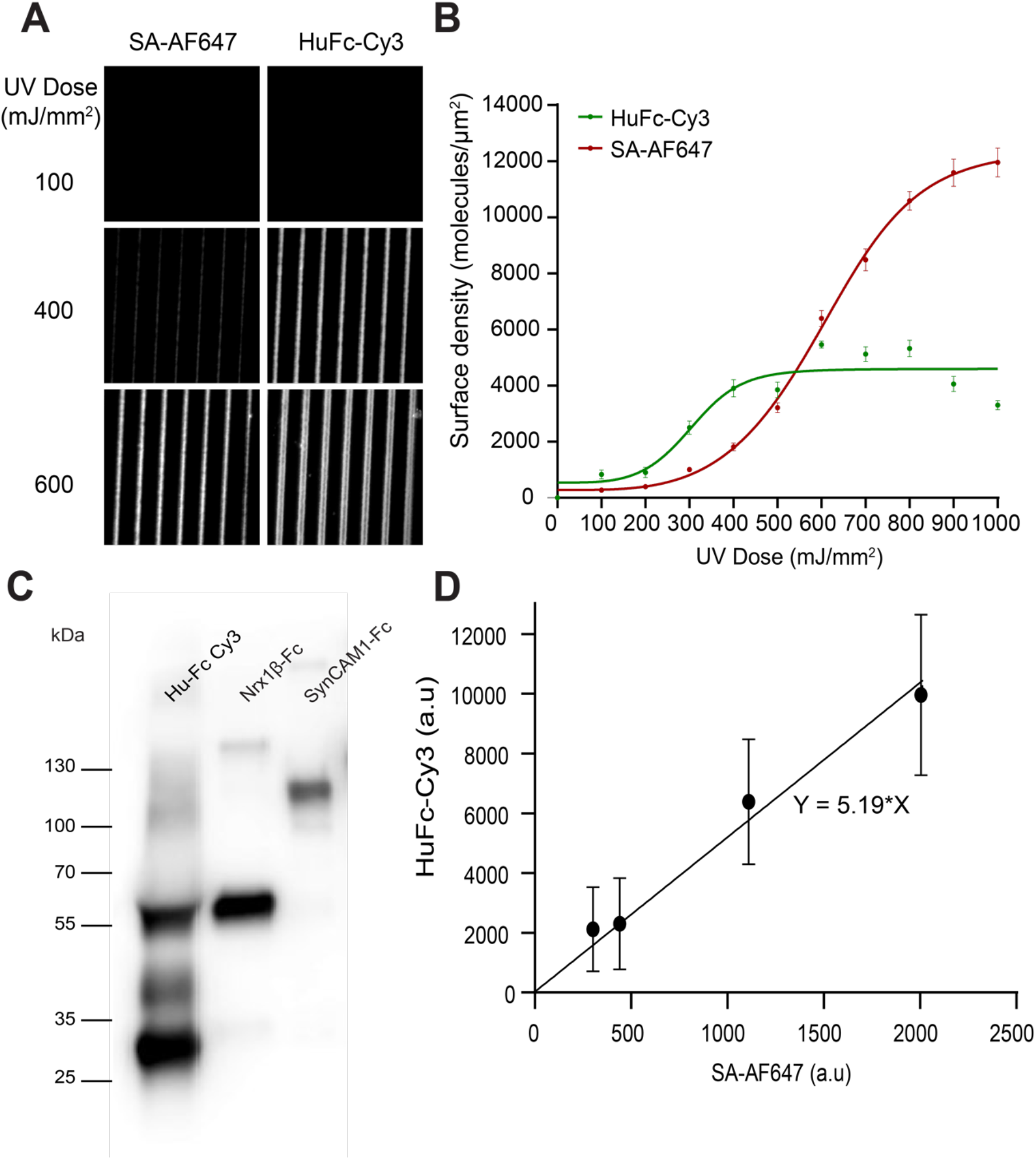
Quantitative micro-photopatterning of synaptic adhesion proteins. (A) Epifluorescence images of the micro-patterned region coated with BSA-biotin and streptavidin-Alexa647, then with biotinylated anti-Fc antibody and Fc-Cy3. (B) Relationship between UV laser dose and fluorescence intensity of micropatterned streptavidin-A647 and Fc-Cy3. Note the saturation in grafted Fc-Cy3 at 600 mJ/mm^2^. (C) Anti-Fc immunoblot of the various purified Fc-tagged proteins used in this study (Nrx1β, SynCAM1, and human Fc alone as a control). (D) Relationship between the fluorescence intensity of micropatterned streptavidin-A647 and the intensity of Fc-Cy3 on the same micro-pattern.

**Figure S2.**
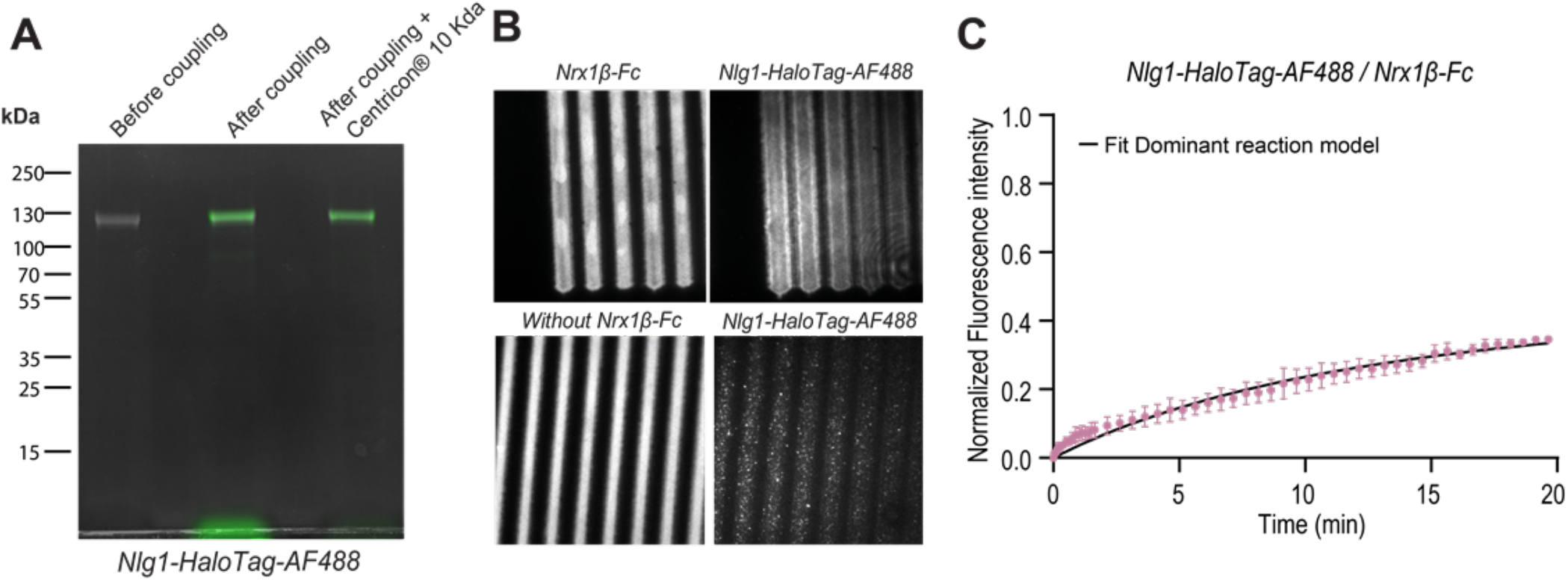
FRAP experiments on soluble Nlg1-HaloTag accumulated at Nrx1β-Fc patterns. (A, B) Purified Nlg1-Halotag-A488 (green) binds to micropatterned lines coated with dimeric Nrx1β-Fc (red), but not to Fc alone. (C) FRAP curves Nlg1-HaloTag-A488 on Nrx1β-Fc pattern.

**Figure S3.**
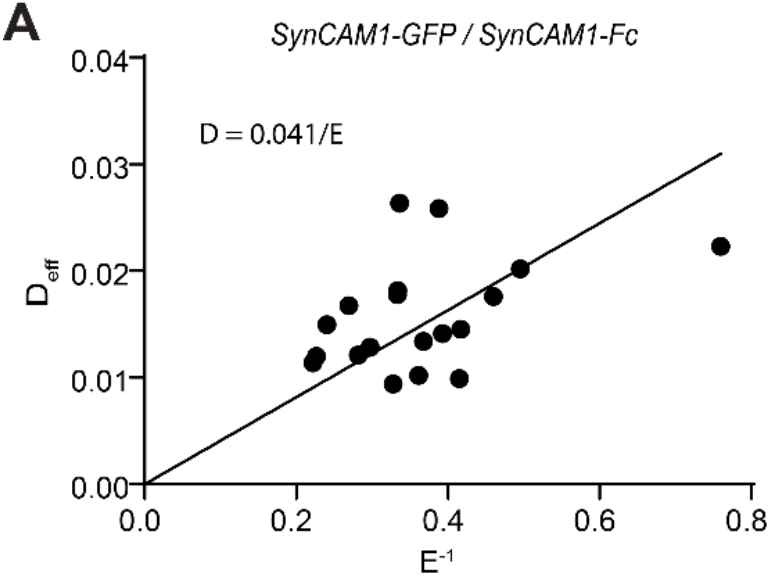
Determination of the coefficient diffusion of free SynCAM1-GFP. (A) Graph showing the correlation between the diffusion coefficient measured by FRAP and the inverse of the enrichment for each individual cell overexpressing SynCAM1-GFP on SynCAM1-Fc patterns.

**Figure S4.**
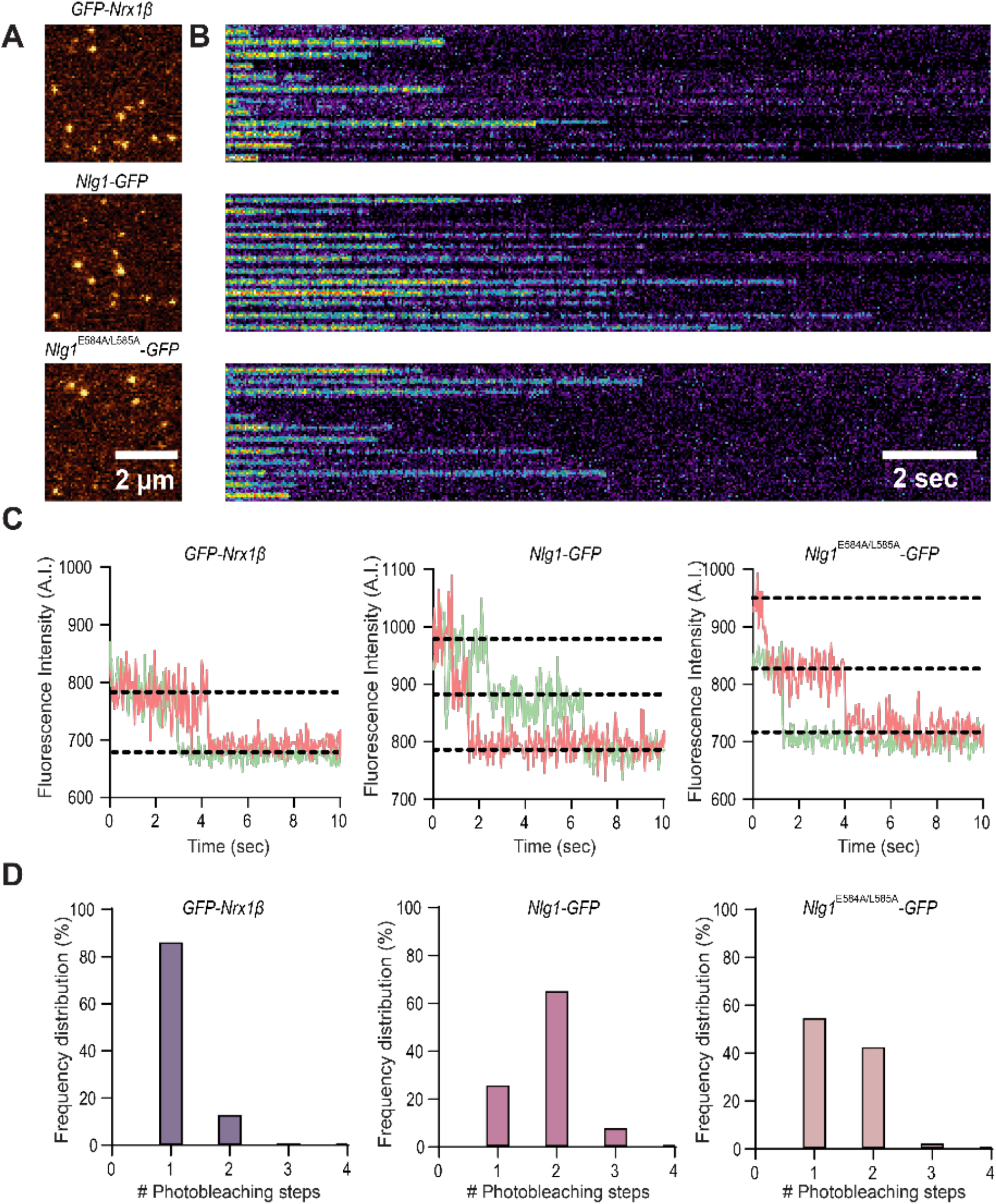
Quantification of the photo-bleaching steps of GFP-Nrx1β and Nlg1-GFP by single molecule pull-down. HEK293T cells expressing GFP-Nrx1β, Nlg1^WT^-GFP, or Nlg1^E584A/L585A^-GFP were lysed and the diluted cell lysates were incubated on glass coverslips coated sequentially with PEG-biotin, neutravidin, and biotinylated anti-GFP antibody, resulting in the binding of individual GFP-tagged membrane molecules. Samples were illuminated with a 491-nm laser in TIRF mode for 20 sec (500 frames, 40 ms exposure), a time after which all molecules were photobleached. (A) Representative images of the first frame of the sequence for the three conditions (GFP-Nrx1β, Nlg1WT-GFP, and Nlg1^E584A/L585A^-GFP). (B) Kymographs showing the fluorescence intensity of 12 independent molecules over time, for each condition. (C) Graphs showing the fluorescence intensity over time for a few individual molecules per condition. Note the sharp photobleaching events that occur as one step (GFP-Nrx1β) or as two steps (for Nlg1-GFP). The Nlg1 ^E584A/L585A^ -GFP mutant exhibits both one-step and two-step photobleaching events. Dashed lines are guides for the eye to show the three fluorescence levels (bottom line = camera background noise, middle line = one GFP molecule, upper line = two GFP molecules). (D) Histograms showing the number of photobleaching steps quantified for 120-130 molecules per condition.

